# Precise label-free quantitative proteomes in high-throughput by microLC and data-independent SWATH acquisition

**DOI:** 10.1101/073478

**Authors:** Jakob Vowinckel, Aleksej Zelezniak, Artur Kibler, Roland Bruderer, Michael Muelleder, Lukas Reiter, Markus Ralser

**Affiliations:** Department of Biochemistry and Cambridge Systems Biology Centre, University of Cambridge, 80 Tennis Court Rd, Cambridge CB2 1GA, UK; The Francis Crick Institute, Mill Hill Laboratory, The Ridgeway, London NW7 1AA, UK; Biognosys AG, Wagistrasse 25, CH8952 Schlieren, Switzerland

## Abstract

While quantitative proteomics is a key technology in biological research, the routine industry and diagnostics application is so far still limited by a moderate throughput, data consistency and robustness. In part, the restrictions emerge in the proteomics dependency on nanolitre/minute flow rate chromatography that enables a high sensitivity, but is difficult to handle on large sample series, and on the stochastic nature in data-dependent acquisition strategies. We here establish and benchmark a label-free, quantitative proteomics platform that uses microlitre/minute flow rate chromatography in combination with data-independent SWATH acquisition. Being able to largely compensate for the loss of sensitivity by exploiting the analytical capacities of microflow chromatography, we show that microLC-SWATH-MS is able to precisely quantify up to 4000 proteins in an hour or less, enables the consistent processing of sample series in high-throughput, and gains quantification precisions comparable to targeted proteomic assays. MicroLC-SWATH-MS can hence routinely process hundreds to thousands of samples to systematically create precise, label free quantitative proteomes.

## INTRODUCTION

Mass spectrometry-based proteomics emerges as prime technology for identifying and quantifying proteins, determining activity, turnover and modification state, and closing gaps in structural biochemistry^1–3^. The routine application of quantitative proteomics in diagnostics^4^ and industrial settings so far hampered by a low throughput (few samples/day), moderate batch-to-batch stability as well as intensive hardware maintenance, all of which render proteomics expensive. Partially, these limits emerge from the proteomics dependency on nanolitre/minute flow rate chromatography (nanoLC) and the necessary electrospray devices (nanospray). Only few years back, nanoLC technology did pave the way for a breakthrough by enabling a level of sensitivity that allows to routinely detect thousands of peptides in a single sample^5^, but the minimal flow rate as well as small capillary and emitter diameters are difficult to handle. Further, nanoLC is relatively slow and susceptible to all sorts of technical distortion^6,7^. Second, on large sample series, further limits arise in stochastic elements of data-dependent acquisition strategies. As these not necessarily quantify each peptide in each replicate or sample, the number of quantifiable proteins becomes inconsistent the larger the sample series becomes.

Some of the original limitations of proteomics do however not necessarily apply any longer, and can be circumvented by the use of new technical developments. First, analytical hardware has substantially progressed recently. Liquid chromatography can now be operated at much higher pressure, allowing higher peak capacity and better resolution in a shorter time. Furthermore, mass spectrometers gained acquisition speed, sensitivity and resolution, so that they can handle faster chromatography and higher dilution rates. Second, data-independent acquisition strategies such as MS^E^ ^8,9^ or SWATH-MS^10,11^ have achieved the necessary precision, so that they routinely overcome the stochastic elements in data acquisition. In parallel, a decade of proteome research has revealed that the typical cellular response involves multiple proteins, which enables identifying biological responses without the need for full proteome coverage, powerful machine learning detects new biological patterns if applied to large sample series^12–14^. As typical proteomic responses can be, in quantitative terms, moderate (fold changes of 1-3x being typical), these approaches however require a high quantitative precision in the datasets. For several applications, the ultimate goal of achieving full proteome coverage has hence secondary importance compared to the imminent need for achieving high quantitative precision and throughout, so that small concentration changes are reliably detected also in large sample series.

A potential solution for key limits in proteomics is capillary, or microflow, chromatography. MicroLC-MS is by principle less sensitive than nanoLC based proteomics, as it operates at 1-2 orders of magnitude higher flow rate than the proteomic-typical nanoLC chromatography. However, the higher flow rate renders microLC compatible with robust analytical-scale electrospray technologies that also allow a high level of control on ionization parameters. Furthermore, by using larger capillary-type columns and electrospray emitter diameters, microLC has shorter dead times, is easier to handle, is more robust, provides a high run-time stability allowing a sophisticated chromatography. Here, we optimize a proteomic set-up which combines microlitre/minute flow rate chromatography (microLC) with data-independent SWATH acquisition, and test its suitability to complement classics set-ups for high-throughput proteomic applications. We find that by exploiting the larger sample capacities of microLC, the reduced sensitivity of microLC compared to nanoLC, can for a large extent be compensated. Preserving the technical advantages of a higher flow rate however, high quantitative precision performance characteristics are achieved in a combination with sequential windowed acquisition of all mass transitions (SWATH-MS)^15^, which enables to overcome the undersampling problem of the fast chromatography. Further, we present a computational batch correction strategy applicable to the quantitative proteome values, so that the routine processing of hundreds to thousands of quantitative proteomes is enabled.

## RESULTS

To find an optimal compromise of flow rate and sensitivity for microLC proteomics, we first evaluated the relationship on a combined nanoLC and microLC chromatographic set-up, coupled to a mass spectrometer selected for a high (up to 100 Hz) acquisition speed so that fast chromatography can to be handled (TripleTOF5600^16^). Signal intensities were determined for standardized peptides (iRT standards^17^) covering flow rates from 300 nL/min to 10 µL/min, using 75 µm (0.3-0.7 µl/min) or 300 µm (1-10 µl/min) inner diameter columns. The 30-fold stepwise increase in flow rate reduced signal intensities by a factor of seven (Fig. 1A). A good compromise between signal and chromatographic quality was found at flow rates between 3 and 5 µL/min (Fig. 1B), and at 3 µL/min, signals were only reduced by a factor of 3.5 compared to nanoLC-MS (Fig. 1A, dotted lines). This decline in signal could be compensated by the higher capacity of the microLC columns, which allow loading of up to 15 µg whole-proteome tryptic digest (around 10x the amount that can be separated in nanoflow chromatography), while still allowing detection of >1200 proteins with as little as 2 µg digest (Fig. 1C). MicroLC-SWATH MS hence requires more samples injection as conventional proteomic set-ups based on nanoLC. Despite a sample quantities represent a common limitation in proteomics, this particular requirement of micro LC represents however a problem only in a small subset of proteomic cases: the typical proteomic sample preparation methods yield ∼10-50 µL of digest, of which in nanoLC-MS/MS typically 1 µL to 2 µL are injected. In microLC-MS-MS, simply a higher injection volume of the same samples is used.

**Figure 1.**
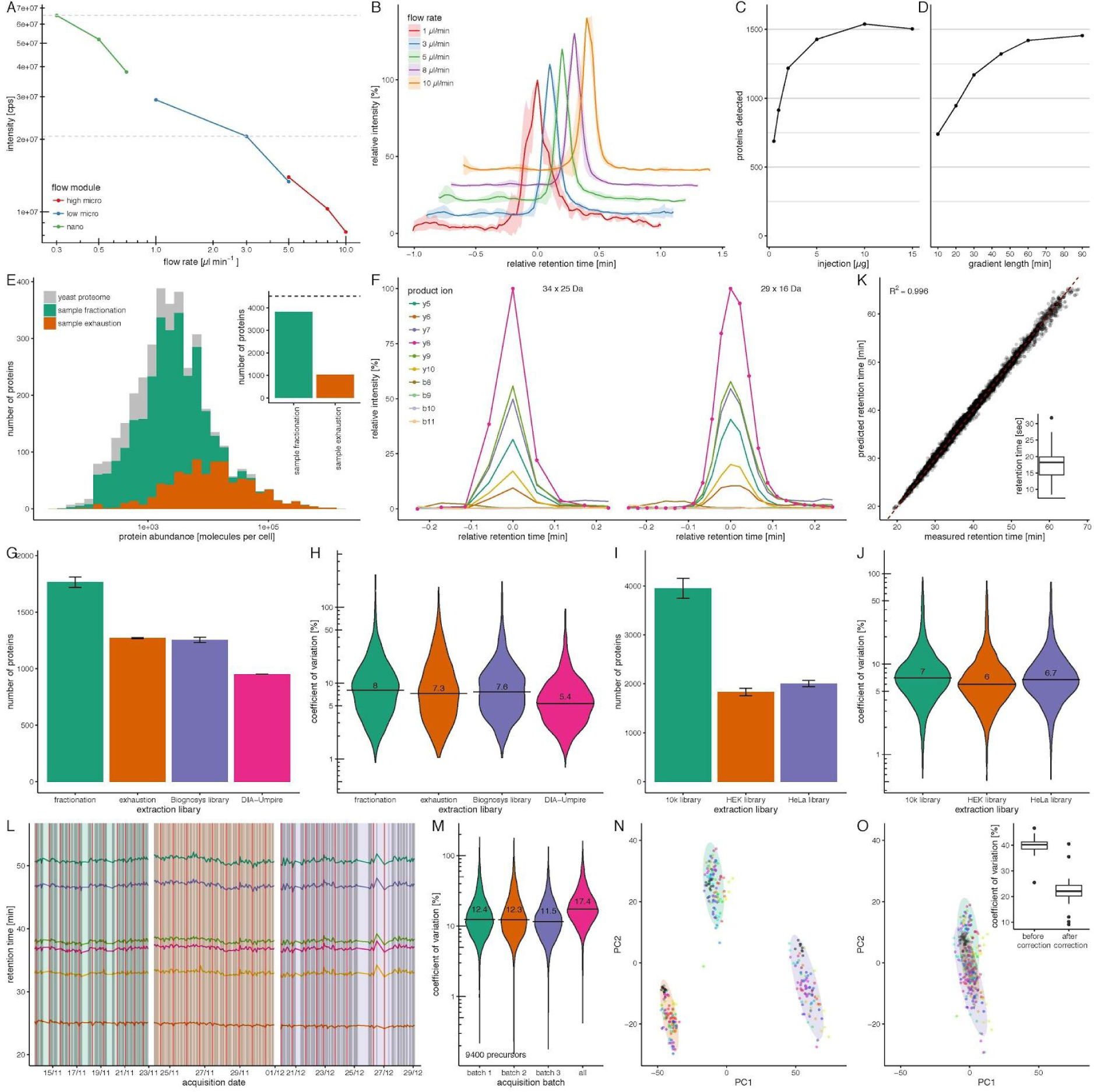
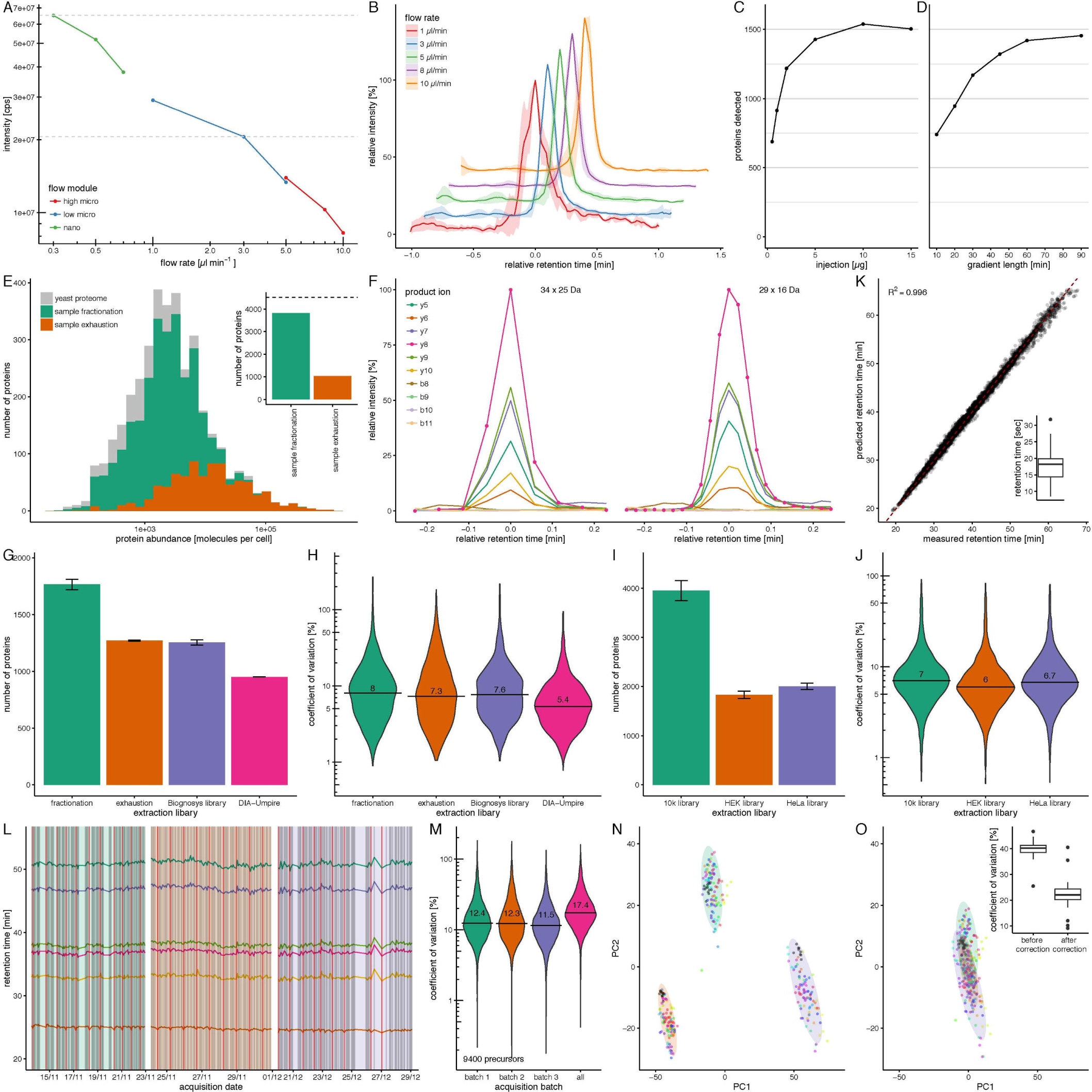
**A. Dependency of signal intensity on flow rate in a proteomic experiment** Combined intensities of standardized peptides (iRT) were determined using nano, low micro, and high-micro flow regimes on an Eksigent 425 system equipped with three respective flow modules and recorded on a TripleTOF5600 mass spectrometer. Signal intensity is a function of the dilution rate, with a factor of 0.3 between 0.3 µl/min and 3 µl/min. **B. Peak characteristics on a microLC set-up** Average precursor peak shapes of 5 iRT peptides determined using flow rates of 1 - 10 µl/min on an Eksigent 3C18-CL-120 column. Chromatography is stable and reproducible in flow rates >1 µl/min. Shaded areas represent standard deviation of signal intensity. **C. 2 µg tryptic digested protein is sufficient to quantify >1200 yeast proteins in microLC-SWATH-MS.** 1-15 µg of yeast protein tryptic digest were injected and separated using a 60 min water to acetonitrile gradient at a flow-rate of 3 µL/min. 914 proteins were identified with 1 µg, 1219 proteins with 2 µg, 1428 proteins with 5 µg and 1504 µg with 15 µg digested protein. Analysis was conducted in Spectronaut v8.0, using a library generated according to Schubert et al.^19^ from a fractionated yeast sample. **D. A 30 min LC gradient is sufficient to quantify >1000 proteins in microLC-SWATH-MS.** 5 µg of yeast protein tryptic digest were injected and separated using water-to-acetonitrile chromatographic gradients of 10-90 min at a flow-rate of 3 µL/min. Extraction of the SWATH spectra yielded quantifiable peptides for 740 proteins (10 min), 946 proteins (20 min), 1170 proteins (30 min), 1322 proteins (45 min), 1420 proteins (60 min) and 1455 proteins (90 min). Analysis conducted in Spectronaut v8.0, using a library generated according to Schubert et al.^19^ from a fractionated yeast sample. **E. SWATH spectral library generation using sample fractionation or exhaustion.** In the sample fractionation approach, a yeast tryptic digest was first separated by high pH reverse phase chromatography on an analytical HPLC as described in methods, and then analyzed in DDA mode with m/z (gas phase) fractionation at 3 µl/min flow rate. In the sample exhaustion approach, a yeast tryptic digest was injected repeatedly until protein identification was saturated. When comparing the proteins identified in both approaches with the published abundances of yeast proteins^18^, the most abundant proteins were identified in both approaches, while proteins with low expression levels were only identified using fractionation. Inset: In total, 3822 (84 %) or 1037 (23 %) out of 4517 expressed yeast proteins^18^ were identified using either method, respectively. **F. Peak representation using microLC-SWATH-MS.** To illustrate peak coverage, an extracted ion chromatogram (XIC) of the peptide TPVITGAPYYER recorded in microLC-SWATH mode using either 34 x 25 m/z or 29 x 16 m/z windows is shown. NanoLC-optimized SWATH^15^ of 34 x 25 m/z with a cycling time of 3.3 s leads to a coverage of 5 points per peak. When limiting the mass range covered to 400-850 m/z and reducing accumulation time to 40 ms, cycling time is 1.32 s and peaks are covered by 11 data points, while being able to capture precursors for 96 % of proteins. **G. Protein quantification in microLC-SWATH-MS using different strategies to construct spectral libraries.** A yeast tryptic digest was analyzed using microLC (0.3 mm x 250 mm Triart-C18, 3 µl/min, 60 min gradient) SWATH-MS by repeated injection of 10 µg digest (9x). Data was processed with Spectronaut v8.0 using SWATH libraries generated by either sample fractionation, sample exhaustion (matrix-matched library), using a spectral library recorded in an unrelated lab (Spectronaut repository), or with a library generated by DIA-Umpire out of the SWATH traces without physically recording a separate spectral library. Data analysis on the basis of the fractionation allowed quantification of 1766 proteins, while 1271 proteins were quantified on the basis of the exhaustion library. The unrelated SWATH library quantified 1256 proteins, and DIA-Umpire 952 proteins. All libraries except the unrelated one (obtained from Spectronaut repository) were generated according to Schubert et al.^19^. **H. Technical variability of yeast protein quantification is low in microLC-SWATH-MS irrespective of data extraction** Fold change (random reference) variability of 677 proteins present in all data sets was compared throughout the nine replicates. Median coefficients of variation are between 7.3 % and 8 % for libraries generated using respectively fractionation and exhaustion approach, 7.6 % for an unrelated yeast library, and 5.4 % for a library generated by DIA-Umpire. All libraries except of obtained from Spectronaut repository were generated according to Schubert et al.^19^ protocol. **I. Human protein quantification using microLC-SWATH** A tryptic digest of a whole-cell protein extract from human K562 cells was analyzed using microLC (0.3 mm x 250 mm Triart-C18, 3 µl/min, 60 min gradient) and coupled to a TripleTOF5600 MS operating in SWATH mode by repeated injection of 3 µg digest (6x). Data was processed with Spectronaut v8.0 using a SWATH library obtained from the SWATHAtlas repository^24^, or using SWATH libraries generated by repeated analysis of HEK293 or HeLa cell extracts (Spectronaut repository). Data analysis using a rich library allows quantification of 4169 proteins, while 2031 proteins can be quantified using a HEK293 and 1906 using a HeLa library, respectively. **J. Technical variability of human protein quantification is low in microLC-SWATH-MS irrespective of data extraction** Fold change variability (random reference) of 726 proteins present in all data sets was compared throughout the six replicates. Median coefficients of variation are around 7 % for all libraries. **K. Retention time stability microLC-SWATH-MS over 327 runs** Correlation between measured apex retention time and predicted retention time. Shown is a representative yeast sample acquired in SWATH mode. Inset: Mean retention time standard deviation of 6 iRT peptides across 327 injections is 17.7 s. **L. Retention times are very stable in microLC-SWATH-MS.** 327 yeast tryptic digest samples spiked with iRT peptides were analyzed on a microLC-SWATH-MS (nanoACQUITY/TripleTOF5600) system in three batches over 3 x 9 days (net acquisition time 16 days, grey vertical lines). Retention times of iRT peptides are shown over time (colored lines), and retention time coefficient of variation for all peptides is lower than 2 % over the whole period. Across the experiment, a control sample was injected repeatedly (red vertical lines) as quality control. **M. Technical variability in microLC-SWATH-MS is low for replicates acquired over a time period of 27 days.** Quality control samples described in L. were analyzed using Spectronaut v8.0, and coefficient of variation for fold changes of 8686 peptides was calculated in batch 1 (green), batch 2 (orange), batch 3 (purple) or across batches (magenta). Intra-batch CVs were around 12 %, while variability over the entire 27 day period was 17.4 %. **N. Variability of 327 yeast proteomes before batch correction.** 38 yeast strains were grown in three batches, and each batch was acquired as three technical replicates in SWATH-MS together with 10-12 evenly interspersed quality control samples. In a PCA, proteomes cluster according to the acquisition batch, with color-coded technical replicates clustering together. **O. Variability of 327 yeast proteomes after batch correction.** After batch correction based on the combined quality control sample profiles, clustering according to batches is reduced, and proteomes cluster according to the color-coded yeast strain. Inset: Median coefficient of variation of peptide intensities between all replicates (technical and biological, 9 replicates per strain) are 39.7 ± 3.2 before batch correction and 22.3 ± 5.4 after batch correction.

In combination with conventional shotgun data-dependent acquisition (DDA), protein identification capacity increased with gradient length, so that >30 minute microflow gradients were sufficient to identify >1000 proteins in a single injection (Fig. 1D). In combination with high pH reversed phase chromatography prefractionation, this set-up did identify 85 % of all expressed yeast open reading frames as detected previously^18^. The overlapping set of 3822 proteins were identified with some abundance bias against the low concentrated proteins (Fig. 1E).

Undersampling is a typical problem in data-dependent acquisition (DDA) strategies. Sharper and better separated chromatographic peaks are analytically desirable, however, fast chromatography amplifies the problem of undersampling. To establish a precise quantitative workflow, we therefore combined microLC with SWATH-MS acquisition, a data-independent acquisition (DIA) strategy where all precursors falling into an isolation window are fragmented simultaneously, and chromatograms are reconstructed computationally post-acquisition^15^. Cycle times of conventional SWATH-MS are in the range of 3 seconds and hence in microLC result in only 4-6 data points covering a peak with a typical 12 s full width at half maximum (FWHM), which would fall short for achieving a precise quantitative dataset. The SWATH regime was hence adapted towards microLC by reducing the windowed dwell time to 40 ms (which on the high concentrated samples was achieved without notable loss in signal and identification capacity (Suppl. Fig. 1)) and limiting segmented acquisition to the precursor-rich mass range between 400-850 m/z. While 85 % of precursors enabling to quantify 96 % of proteins fall into this mass range (Suppl. Fig. 2), the modification allowed shortening the cycle time to 1.3s, which enabled to cover a typical microLC chromatographic peak with a critical amount of 8-12 data points (Fig. 1F).

On this setup we acquired replicate injections of 10 µg of a whole-cell yeast digest using 60 min gradients and 3 µl/min, and for comparison, used different strategies for SWATH-MS data extraction. SWATH spectral libraries were generated using proteome prefractionation (following original approaches^15,19^), repeated injection of a sample mixture matching the actual sample matrix (‘exhaustion’), or by pseudo-MS/MS correlative precursor-fragment feature extraction using DIA-Umpire^20^. All libraries were generated using the same protocol^19^ at <1 % FDR using a combination of X! Tandem^21^ and Comet^22^ search engines. Extracting a typical microLC-SWATH run using the spectral library created by prefractionation, we could quantify 1766±46 yeast proteins using 34x25 m/z SWATH windows, or 1422±53 proteins when using 29x16 m/z windows, respectively (Fig. 1G and Suppl. Fig. 3). Analysis of the same data using the library generated by exhaustion allowed quantification of 1271±5 and 1157±13 proteins, which was similar to a public library generated by nanoLC-MS/MS (Spectronaut^23^ repository) yielding 1256±23 and 1118±26 proteins, respectively. In the absence of separately acquired spectral library, DIA-Umpire instead did quantify 952±0 and 890±2 proteins (Fig. 1G and Suppl. Fig. 3). Also in terms of peptide quantification, peak reconstruction on the basis of a prefractionation-based library performed best, followed by the public and exhaustion-based libraries (Suppl. Fig. 4 and Suppl. Fig. 5). Next, we tested the performance of microLC-SWATH-MS on a standardized whole-proteome human cell line (K562) digest, by extracting data using three publicly available spectral libraries generated by combining multiple tissues and fractionation^24^ or by repetitive injection of tissue-specific cell digests of HEK293 or HeLa cells (Spectronaut^23^ repository). MicroLC-SWATH-MS achieved quantification of 3951±205, 1832±74 and 2007±63 proteins, respectively, out of single-injections of the unfractionated K562 digest, with peptide numbers following the same trend (Fig. 1I, Suppl. Fig. 7).

The MicroLC-SWATH-MS optimized in this way did yield precise quantities in label free proteomics. Median coefficients of variation (CVs) for replicate injections in all acquisition strategies and analysis libraries were in the range of 5.4-8.8 % for both peak areas as well as protein fold changes for the yeast samples (Fig. 1H and Suppl. Fig. 6), and 5.5-7 % for the standardized human sample. The precision was similar both for low and high concentrated peptides, over the full range of five orders of magnitude (Fig. 1J, Suppl. Fig. 8, Suppl. Fig. 9 and Suppl. Fig. 10). Of note, on our dataset, the library generated with DIA-Umpire quantified less proteins, but yielded the best precision values (Fig 1H). Being label-free, microLC-SWATH-MS strategies thus clearly equal or even exceeds, precision values as typically obtained in a label-free quantitative proteomic experiment^15,25–27^.

Applied to large sample series that are enabled by the fast throughput of microLC-SWATH-MS, such high precision values facilitate to detect small and moderate protein concentration changes, that with a technology of low precision would be lost due to error propagation. On large proteomic sample series additional limits emerge however from unavoidable batch effects. If left uncorrected, these can exceed that of the biological information^28,29^. To monitor batch characteristics and to develop counter-measures, we analyzed 296 *Saccharomyces cerevisiae* proteomes (38 yeast cultures grown in nine replicates, 3 biological x 3 technical) and acquired them in three batches. We repetitively included a mixture of all samples as quality control (QC) sample every 10-12 injections, which in total resulted in 327 whole-proteome samples processed (Fig. 1L). MicroLC was, even without retention time normalization, highly stable (standard deviation of apex retention times 17.7 s in 60 min gradients over the 327 runs, Fig. 1K, inset) both within as well as across batches (Fig. 1L). Within the batches spanning 100-116 whole proteome digests each acquired over a period of around 9 days (net acquisition time 6 days), median CVs of QC samples were around 12 %, while QC CVs between batches were 17.4 % (Fig. 1M). Correction for the chromatographic drifts was effectively accomplished by the use of retention time standards and linear normalisation^17^. To address the more complicated batch effects that emerge from signal intensity drifts and fluctuations in the ion traces, we adapted a strategy based on models developed for gene expression analysis^28^ making use of the QC samples to monitor instrument performance. Intensity correction reduced signal variability by 43 % that largely correspond to the batch effect, while maintaining each individual strain’s proteome profile (Fig. 1N, Fig. 1O, Suppl. Fig. 11 and Suppl. Fig. 12). Applying these strategies to the 327 whole proteomes yielded the quantification of 1212±100 proteins in each sample, with the average peptide quantified with CV of 22.3±5.4 % across all technical and biological replicates.

In conclusion, we show that microLC-MS/MS in combination with sample fractionation is able to achieve competitive protein identification characteristics to that of canonical proteomics platforms, while in combination with data independent acquisition and SWATH-MS, the platform can capture a quantitative snapshot of a yeast or human proteome in an hour or less and in high throughput. Achieving high precision and signal stability in label free proteomics, microLC-SWATH-MS enabled processing hundreds of samples in series and in a few days. The performance characteristics are illustrated by quantifying 1210±100 proteins across 327 whole proteomes, achieved in a net acquisition time of solely 17 days, and with CV values that are comparable to that of FDA approved targeted assays. The technology is hence ideal for large proteomic sample series, and for applications where precision and quantitative robustness are the key objectives. This includes, for instance, the application of machine learning strategies to detect novel biological patterns, for the analysis of biological time series, and for comparative studies in basic research and diagnostics that require large sample series. Indeed, the high flow rate renders microLC-SWATH-MS a highly robust proteomic technology, with low instrument downtime and maintenance cycles, so it is ideal for laboratories that address large sample number in diagnostics and for data-driven biology.

## Acknowledgments

We are glad to the Francis Crick Institute, the Wellcome Trust (RG 093735/Z/10/Z) and the ERC (Starting grant 260809) for funding. A.Z. is EMBO fellow (ALTF-969 2014) which is co-funded by the European Commission (LTFCOFUND2013, GA-2013-609409) support from Marie Curie Actions.

## Competitive Interest statement

The authors J.V., R.B. and L.R. are employees of Biognosys AG (Switzerland). Spectronaut is a trademark of Biognosys AG.

## Materials and Methods

### Materials, solutions and reagents

Chemicals and reagents were obtained from Sigma unless stated otherwise, and UPLC/MS grade chromatographic solvents were from Greyhound.

### Sample preparation for mass spectrometry

A standardized yeast sample was generated by growing the prototrophic *S. cerevisiae* strain YSBN1^30^ in Yeast Nitrogen Base (YNB) medium without amino acids containing 2 % glucose until mid-exponential phase. Cells were harvested by centrifugation and snap-frozen in aliquots equaling 10 OD. Sample preparation was performed based^31^ with the following modifications. Cells were broken by bead shaking with 200 µl 0.05 M ammonium bicarbonate in a FastPrep instrument (3x 30 s, 6.5 m/s, 4 °C), and cell pellet after centrifugation was re-extracted with 200 µl lysis buffer (0.1 M NaOH, 0.05 M EDTA, 2 % SDS, 2 % 2-mercaptoethanol) for 10 min at 90 °C, and again for 10 min at 90 °C after addition of 0.1 M acetic acid. Combined supernatants were precipitated using 10 % TCA, and processed further according to the RapiGest protocol^31^. Protein concentration before digest was determined using Pierce BCA assay kit (Thermo), and adjusted to 2 µg/µl with 0.2 % RapiGest SF (Waters) in ABC. MS compatible human protein digest from K562 cells was obtained from Promega.

For library generation using pre-fractionation, 1 mg of yeast digest was separated by high pH reverse phase chromatography on a Waters ACQUITY instrument. A reverse phase column (Waters, BEH C18, 2.1 × 150 mm, 1.7 µm) was utilized in combination with a 20 mM ammonium formate to 20 mM ammonium formate/ 80 % ACN gradient, collecting 33 fractions. Before analysis, samples were spiked with 0.5x HRM kit (Biognosys). QC samples were prepared as a mixture of 10 % of each individual sample.

### Chromatography

Chromatographic separation was performed either on an Ekspert NanoLC 425 system (SCIEX) for combined nano and micro flow analysis, or a nanoACQUITY system (Waters) for microflow-only sample series. In nano flow, the NanoLC 425 system was equipped with a nano flow module, and samples were first loaded onto a trap column (Chrom XP C18-3µm, 0.12 nm, 0.35 x 0.5 mm) by isocratically running the system at a flow rate of 5 µl/min for 6 min with 0.1 % formic acid (FA) in water. Peptides were then eluted onto the analytical column (3C18-CL-120, 3 µm, 0.12 nm, 0.075 x 150 mm, Eksigent) and separated on a linear gradient of 2-30 % 0.1 % FA in acetonitrile (ACN) in 25 min. For micro flow analysis, the same system was equipped with a low micro flow module (1-5 µl/min) or high micro flow module (5-10 µl/min) and set up for direct injection onto an analytical column (3C18-CL-120, 3 µm, 120 Å, 0.3 x 150 mm, Eksigent). Separation was performed on a linear gradient of 2-30 % 0.1 % FA in ACN in 25 min. For micro flow on the nanoACQUITY system, the sample manager was set up in direct injection mode and equipped with a Triart C18 column (0.12 nm, 3 µm, 0.3 mm x 250 mm, YMC). After injecting samples onto the analytical column, peptides were separated on linear gradients detailed in Suppl. table 1.

### Mass Spectrometry

Analysis was performed on a Tandem Quadrupole Time-of-Flight mass spectrometer (SCIEX TripleTOF5600^16^) coupled to either a Nanospray III Ion Source (SCIEX) or a DuoSpray Ion Source (SCIEX), controlled by Analyst software (v.1.6). For nano flow, the ion source was equipped with 10 µm SilicaTip electrospray emitters (New Objective) and parameters were as described^31^. For micro flow, the ion source was equipped with a 25 µm TurboIonSpray probe (SCIEX), and parameters were as follows: ISVF = 5500, GS1 = 10, GS2 = 0, CUR = 25, TEM = 100.

To acquire spectral libraries used in SWATH data extraction, the mass spectrometer was operated in information-dependent acquisition (IDA) and high sensitivity mode, with first a 250 ms TOF MS survey scan over a mass range of 400-1250 m/z, followed by 100 ms MS/MS scans of 20 ion candidates per cycle with dynamic background subtraction. The selection criteria for the parent ions included the intensity, where ions had to be greater than 150 cps, with a charge state between 2 and 4. The dynamic exclusion duration was set for 1 s. Collision-induced dissociation was triggered by rolling collision energy. For generation of SWATH libraries by sample fractionation, precursor-rich fractions were further injected twice, with gas phase fractionation of 400-650 and 650-1250. For data-independent acquisition, the instrument was operated in SWATH mode with selection windows detailed in Suppl. table 2 for mass ranges of 400-1250 or 400-850 m/z. With accumulation times of 100 and 40 ms cycle times were 3.3 s and 1.2 s, respectively.

### Data analysis and SWATH library generation

All SWATH assay libraries were built following Schubert et al^19^. Briefly, spectral data acquired in IDA mode was centroided using qtofpeakpicker^32^. Centroided files were searched with X! Tandem^21^ and Comet^22^ against annotated yeast proteins database with included reversed decoy peptides. Search results were scored using PeptideProphet and combined with iProphet. Mayu^33^ was used to estimate iProphet probabilities to control for protein identification false discovery rate (FDR <1 %). The final spectral library was assembled using SpectraST^34^ by retaining spectra above iProphet FDR controlled cutoff and normalizing chromatography to iRT peptide retention time reference. SpectraST output was then converted to tsv format suitable for Spectronaut retaining 6 most intense transitions of y and b ions using spectrast2tsv from msprotoemicstools^35^. For large-scale data analysis of 327 yeast samples, a minimal consensus library constructed from exhaustion-based IDA acquisitions was compiled in Spectronaut. In the DIA-Umpire approach, we extracted precursor-fragment features by applying signal extraction module from DIA-Umpire workflow using default recommended parameters for TripleTOF5600 instrument on data acquired in SWATH mode by sample exhaustion of the yeast proteome. Generated pseudo MS/MS mgf files were converted into mzXML and further subjected to database search and processed as described above to generate SWATH assay library. All SWATH data quantification was performed in Spectronaut (v. 8.0.9600, Biognosys) using default settings. Publicly available SWATH libraries used were obtained from the Biognosys library repository or from SWATHAtlas^24^.

For visualization of chromatographic peaks, data of selected peptides was analyzed in Skyline^36^ (v. 3.5.0.9191) with SWATH isolation windows detailed in Suppl. table 2, and chromatograms of precursors and products exported as text files. Post-processing was conducted in R^37^ by first removing precursors from all samples where the median Qvalue was > 0.01, and then transforming all remaining precursors with Qvalue > 0.01 into NA. Injection differences were corrected by a robust sum approach, and peptides belonging to one protein were selected based on closest fold change correlation. To account for confounding effects related to acquisition dates, we performed batch correction by introducing QC samples in experimental design. External standard QC samples were prepared as a mixture of all injected samples and were measured every 10-12 samples. Each MS acquisition batch had >10 QC samples allowing to correct for the most evident batch effects attributed to an acquisition date (Suppl. Fig. 11). Signal correction was performed using ComBat approach^38^ as implemented in R sva package^39^. Plotting was performed in ggplot2 package^40^.

## References

1 Walzthoeni, T., Leitner, A., Stengel, F. & Aebersold, R. Mass spectrometry supported determination of protein complex structure. Curr. Opin. Struct. Biol. 23, 252–260 (2013).

2 Angel, T. E. et al. Mass spectrometry-based proteomics: existing capabilities and future directions. Chem. Soc. Rev. 41, 3912–3928 (2012).

3 Cardoza, J. D., Parikh, J. R., Ficarro, S. B. & Marto, J. A. Mass spectrometry-based proteomics: qualitative identification to activity-based protein profiling. Wiley Interdiscip. Rev. Syst. Biol. Med. 4, 141–162 (2012).

4 Muntel, J. et al. Advancing Urinary Protein Biomarker Discovery by Data-Independent Acquisition on a Quadrupole-Orbitrap Mass Spectrometer. J. Proteome Res. 14, 4752–4762 (2015).

5 Valaskovic, G. A. & Kelleher, N. L. Miniaturized formats for efficient mass spectrometry-based proteomics and therapeutic development. Curr. Top. Med. Chem. 2, 1–12 (2002).

6 Mitulović, G. & Mechtler, K. HPLC techniques for proteomics analysis—a short overview of latest developments. Brief. Funct. Genomic. Proteomic. 5, 249–260 (2006).

7 Wilm, M. & Mann, M. Analytical properties of the nanoelectrospray ion source. Anal. Chem. 68, 1–8 (1996).

8 Kustatscher, G. et al. Proteomics of a fuzzy organelle: interphase chromatin. EMBO J. 33, 648–664 (2014).

9 Soste, M. et al. A sentinel protein assay for simultaneously quantifying cellular processes. Nat. Methods 11, 1045–1048 (2014).

10 Gillet, L. C. et al. Targeted data extraction of the MS/MS spectra generated by data-independent acquisition: a new concept for consistent and accurate proteome analysis. Mol. Cell. Proteomics 11, O111.016717 (2012).

11 Andrews, G. L., Simons, B. L., Young, J. B., Hawkridge, A. M. & Muddiman, D.C. Performance characteristics of a new hybrid quadrupole time-of-flight tandem mass spectrometer (TripleTOF 5600). Anal. Chem. 83, 5442–5446 (2011).

12 Escher, C. et al. Using iRT, a normalized retention time for more targeted measurement of peptides. Proteomics 12, 1111–1121 (2012).

13 Ghaemmaghami, S. et al. Global analysis of protein expression in yeast. Nature 425, 737–741 (2003).

14 Schubert, O. T. et al. Building high-quality assay libraries for targeted analysis of SWATH MS data. Nat. Protoc. 10, 426–441 (2015).

15 Tsou, C.-C. et al. DIA-Umpire: comprehensive computational framework for data-independent acquisition proteomics. Nat. Methods 12, 258–64, 7 p following 264 (2015).

16 Craig, R. & Beavis, R. C. TANDEM: matching proteins with tandem mass spectra. Bioinformatics 20, 1466–1467 (2004).

17 Eng, J. K., Jahan, T. A. & Hoopmann, M. R. Comet: an open-source MS/MS sequence database search tool. Proteomics 13, 22–24 (2013).

18 Bruderer, R. et al. Extending the limits of quantitative proteome profiling with data-independent acquisition and application to acetaminophen-treated three-dimensional liver microtissues. Mol. Cell. Proteomics 14, 1400–1410 (2015).

19 Rosenberger, G. et al. A repository of assays to quantify 10,000 human proteins by SWATH-MS. Sci Data 1, 140031 (2014).

20 Basak, T., Bhat, A., Malakar, D., Pillai, M. & Sengupta, S. In-depth comparative proteomic analysis of yeast proteome using iTRAQ and SWATH based MS. Mol. Biosyst. 11, 2135–2143 (2015).

21 Selevsek, N. et al. Reproducible and consistent quantification of the Saccharomyces cerevisiae proteome by SWATH-mass spectrometry. Mol. Cell. Proteomics 14, 739–749 (2015).

22 Burniston, J. G., Connolly, J., Kainulainen, H., Britton, S. L. & Koch, L. G. Label-free profiling of skeletal muscle using high-definition mass spectrometry. Proteomics 14, 2339–2344 (2014).

23 Gregori, J. et al. Batch effects correction improves the sensitivity of significance tests in spectral counting-based comparative discovery proteomics. J. Proteomics 75, 3938–3951 (2012).

24 Scherer, A. Batch Effects and Noise in Microarray Experiments: Sources and Solutions. (John Wiley & Sons, 2009).

25 Canelas, A. B. et al. Integrated multilaboratory systems biology reveals differences in protein metabolism between two reference yeast strains. Nat. Commun. 1, 145 (2010).

26 Vowinckel, J. et al. The beauty of being (label)-free: sample preparation methods for SWATH-MS and next-generation targeted proteomics. F1000Res. 2, (2014).

27 Chambers, M. C. et al. A cross-platform toolkit for mass spectrometry and proteomics. Nat. Biotechnol. 30, 918–920 (2012).

28 Reiter, L. et al. Protein identification false discovery rates for very large proteomics data sets generated by tandem mass spectrometry. Mol. Cell. Proteomics 8, 2405–2417 (2009).

29 Deutsch, E. W. et al. A guided tour of the Trans-Proteomic Pipeline. Proteomics 10, 1150–1159 (2010).

30 Aebersold, R. et al. msproteomicstools. at <https://github.com/msproteomicstools>

31 MacLean, B. et al. Skyline: an open source document editor for creating and analyzing targeted proteomics experiments. Bioinformatics 26, 966–968 (2010).

32 R Core Team. R: A Language and Environment for Statistical Computing. (2015). at <https://www.R-project.org/>

33 Johnson, W. E., Li, C. & Rabinovic, A. Adjusting batch effects in microarray expression data using empirical Bayes methods. Biostatistics 8, 118–127 (2007).

34 Leek, J. T., Johnson, W. E., Parker, H. S., Jaffe, A. E. & Storey, J. D. The sva package for removing batch effects and other unwanted variation in high-throughput experiments. Bioinformatics 28, 882–883 (2012).

35 Wickham, H. ggplot2: Elegant Graphics for Data Analysis. (2009). at <http://had.co.nz/ggplot2/book>

## References

(1) Walzthoeni, T.; Leitner, A.; Stengel, F.; Aebersold, R. Curr. Opin. Struct. Biol. 2013, 23 (2), 252–260.

(2) Angel, T. E.; Aryal, U. K.; Hengel, S. M.; Baker, E. S.; Kelly, R. T.; Robinson, E. W.; Smith, R. D. Chem. Soc. Rev. 2012, 41 (10), 3912–3928.

(3) Cardoza, J. D.; Parikh, J. R.; Ficarro, S. B.; Marto, J. A. Wiley Interdiscip. Rev. Syst. Biol. Med. 2012, 4 (2), 141–162.

(4) Muntel, J.; Xuan, Y.; Berger, S. T.; Reiter, L.; Bachur, R.; Kentsis, A.; Steen, H. J. Proteome Res. 2015, 14 (11), 4752–4762.

(5) Valaskovic, G. A.; Kelleher, N. L. Curr. Top. Med. Chem. 2002, 2 (1), 1–12.

(6) Mitulović, G.; Mechtler, K. Brief. Funct. Genomic. Proteomic. 2006, 5 (4), 249–260.

(7) Wilm, M.; Mann, M. Anal. Chem. 1996, 68 (1), 1–8.

(8) Muntel, J.; Fromion, V.; Goelzer, A.; Maass, S.; Mader, U.; Buttner, K.; Hecker, M.; Becher, D. Mol. Cell. Proteomics 2014, 13 (4), 1008–1019.

(9) Plumb, R. S.; Johnson, K. A.; Paul, R.; Smith, B. W.; Wilson, I. D.; Castro-Perez, J. M.; Nicholson, J. K. Rapid Commun. Mass Spectrom. 2006, 20 (13), 1989–1994.

(10) Selevsek, N.; Chang, C.-Y.; Gillet, L. C.; Navarro, P.; Bernhardt, O. M.; Reiter, L.; Cheng, L.-Y.; Vitek, O.; Aebersold, R. Mol. Cell. Proteomics 2015, 14 (3), 739–749.

(11) Gillet, L. C.; Navarro, P.; Tate, S.; Röst, H.; Selevsek, N.; Reiter, L.; Bonner, R.; Aebersold, R. Mol. Cell. Proteomics 2012, 11 (6), O111.016717.

(12) Kustatscher, G.; Hégarat, N.; Wills, K. L. H.; Furlan, C.; Bukowski-Wills, J.-C.; Hochegger, H.; Rappsilber, J. EMBO J. 2014, 33 (6), 648–664.

(13) Soste, M.; Hrabakova, R.; Wanka, S.; Melnik, A.; Boersema, P.; Maiolica, A.; Wernas, T.; Tognetti, M.; von Mering, C.; Picotti, P. Nat. Methods 2014, 11 (10), 1045–1048.

(14) Zou, Q.; Quan, Z. Curr. Proteomics 2016, 13 (2), 77–78.

(15) Gillet, L. C.; Navarro, P.; Tate, S.; Röst, H.; Selevsek, N.; Reiter, L.; Bonner, R.; Aebersold, R. Mol. Cell. Proteomics 2012, 11 (6), O111.016717.

(16) Andrews, G. L.; Simons, B. L.; Young, J. B.; Hawkridge, A. M.; Muddiman, D. C. Anal. Chem. 2011, 83 (13), 5442–5446.

(17) Escher, C.; Reiter, L.; MacLean, B.; Ossola, R.; Herzog, F.; Chilton, J.; MacCoss, M. J.; Rinner, O. Proteomics 2012, 12 (8), 1111–1121.

(18) Ghaemmaghami, S.; Huh, W.-K.; Bower, K.; Howson, R. W.; Belle, A.; Dephoure, N.; O’Shea, E. K.; Weissman, J. S. Nature 2003, 425 (6959), 737–741.

(19) Schubert, O. T.; Gillet, L. C.; Collins, B. C.; Navarro, P.; Rosenberger, G.; Wolski, W. E.; Lam, H.; Amodei, D.; Mallick, P.; MacLean, B.; Aebersold, R. Nat. Protoc. 2015, 10 (3), 426–441.

(20) Tsou, C.-C.; Avtonomov, D.; Larsen, B.; Tucholska, M.; Choi, H.; Gingras, A.-C.; Nesvizhskii, A. I. Nat. Methods 2015, 12 (3), 258–264, 7 p following 264.

(21) Craig, R.; Beavis, R. C. Bioinformatics 2004, 20 (9), 1466–1467.

(22) Eng, J. K.; Jahan, T. A.; Hoopmann, M. R. Proteomics 2013, 13 (1), 22–24.

(23) Bruderer, R.; Bernhardt, O. M.; Gandhi, T.; Miladinović, S. M.; Cheng, L.-Y.; Messner, S.; Ehrenberger, T.; Zanotelli, V.; Butscheid, Y.; Escher, C.; Vitek, O.; Rinner, O.; Reiter, L. Mol. Cell. Proteomics 2015, 14 (5), 1400–1410.

(24) Rosenberger, G.; Koh, C. C.; Guo, T.; Röst, H. L.; Kouvonen, P.; Collins, B. C.; Heusel, M.; Liu, Y.; Caron, E.; Vichalkovski, A.; Faini, M.; Schubert, O. T.; Faridi, P.; Ebhardt, H. A.; Matondo, M.; Lam, H.; Bader, S. L.; Campbell, D. S.; Deutsch, E. W.; Moritz, R. L.; Tate, S.; Aebersold, R. Sci Data 2014, 1, 140031.

(25) Basak, T.; Bhat, A.; Malakar, D.; Pillai, M.; Sengupta, S. Mol. Biosyst. 2015, 11 (8), 2135–2143.

(26) Selevsek, N.; Chang, C.-Y.; Gillet, L. C.; Navarro, P.; Bernhardt, O. M.; Reiter, L.; Cheng, L.-Y.; Vitek, O.; Aebersold, R. Mol. Cell. Proteomics 2015, 14 (3), 739–749.

(27) Burniston, J. G.; Connolly, J.; Kainulainen, H.; Britton, S. L.; Koch, L. G. Proteomics 2014, 14 (20), 2339–2344.

(28) Gregori, J.; Josep, G.; Laura, V.; Olga, M.; Alex, S.; José, B.; Josep, V. J. Proteomics 2012, 75 (13), 3938–3951.

(29) Scherer, A. Batch Effects and Noise in Microarray Experiments: Sources and Solutions; John Wiley & Sons, 2009.

(30) Canelas, A. B.; Harrison, N.; Fazio, A.; Zhang, J.; Pitkänen, J.-P.; Brink, J. van D.; Bakker, B. M.; Bogner, L.; Bouwman, J.; Castrillo, J. I.; Cankorur, A.; Chumnanpuen, P.; Daran-Lapujade, P.; Dikicioglu, D.; Eunen, K. van; Ewald, J. C.; Heijnen, J. J.; Kirdar, B.; Mattila, I.; Mensonides, F. I. C.; Niebel, A.; Penttilä, M.; Pronk, J. T.; Reuss, M.; Salusjärvi, L.; Sauer, U.; Sherman, D.; Siemann-Herzberg, M.; Westerhoff, H.; Winde, J. de; Petranovic, D.; Oliver, S. G.; Workman, C. T.; Zamboni, N.; Nielsen, J. Nat. Commun. 2010, 1, 145.

(31) Vowinckel, J.; Capuano, F.; Campbell, K.; Deery, M. J.; Lilley, K. S.; Ralser, M. F1000Res. 2014, 2.

(32) Chambers, M. C.; Maclean, B.; Burke, R.; Amodei, D.; Ruderman, D. L.; Neumann, S.; Gatto, L.; Fischer, B.; Pratt, B.; Egertson, J.; Hoff, K.; Kessner, D.; Tasman, N.; Shulman, N.; Frewen, B.; Baker, T. A.; Brusniak, M.-Y.; Paulse, C.; Creasy, D.; Flashner, L.; Kani, K.; Moulding, C.; Seymour, S. L.; Nuwaysir, L. M.; Lefebvre, B.; Kuhlmann, F.; Roark, J.; Rainer, P.; Detlev, S.; Hemenway, T.; Huhmer, A.; Langridge, J.; Connolly, B.; Chadick, T.; Holly, K.; Eckels, J.; Deutsch, E. W.; Moritz, R. L.; Katz, J. E.; Agus, D. B.; MacCoss, M.; Tabb, D. L.; Mallick, P. Nat. Biotechnol. 2012, 30 (10), 918–920.

(33) Reiter, L.; Claassen, M.; Schrimpf, S. P.; Jovanovic, M.; Schmidt, A.; Buhmann, J. M.; Hengartner, M. O.; Aebersold, R. Mol. Cell. Proteomics 2009, 8 (11), 2405–2417.

(34) Deutsch, E. W.; Mendoza, L.; Shteynberg, D.; Farrah, T.; Lam, H.; Tasman, N.; Sun, Z.; Nilsson, E.; Pratt, B.; Prazen, B.; Eng, J. K.; Martin, D. B.; Nesvizhskii, A. I.; Aebersold, R. Proteomics 2010, 10 (6), 1150–1159.

(35) Aebersold, R.; Blum, L.; Gillet, L.; Malmström, L.; Navarro, P.; Röst, H.; Rosenberger, G.; Sigurdur, S. msproteomicstools https://github.com/msproteomicstoolshttp://paperpile.com/b/fBTvYQ/ss9Exhttps://github.com/msproteomicstools (accessed Jan 19, 2016).

(36) MacLean, B.; Tomazela, D. M.; Shulman, N.; Chambers, M.; Finney, G. L.; Frewen, B.; Kern, R.; Tabb, D. L.; Liebler, D. C.; MacCoss, M. J. Bioinformatics 2010, 26 (7), 966–968.

(37) R Core Team. R Foundation for Statistical Computing: Vienna, Austria 2015.

(38) Johnson, W. E.; Li, C.; Rabinovic, A. Biostatistics 2007, 8 (1), 118–127.

(39) Leek, J. T.; Johnson, W. E.; Parker, H. S.; Jaffe, A. E.; Storey, J. D. Bioinformatics 2012, 28 (6), 882–883.

(40) Wickham, H. Springer-Verlag New York 2009.

